## Supplementary figures for "Beyond benchmarking: towards predictive models of dataset-specific single-cell RNA-seq pipeline performance"

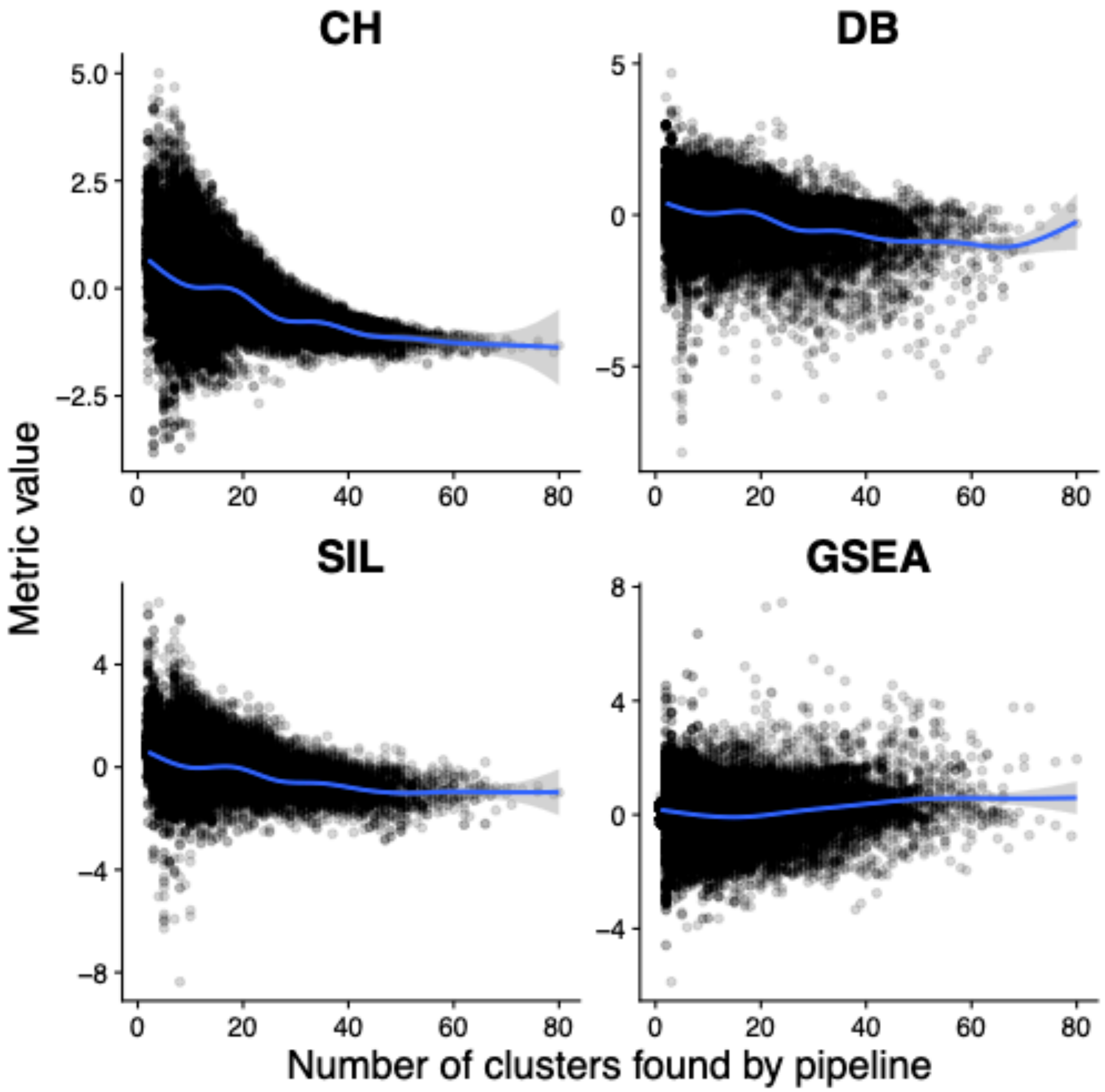


**Supplementary Figure 1** Correlation of the four unsupervised metrics with the number of clusters found by a given pipeline across all datasets.

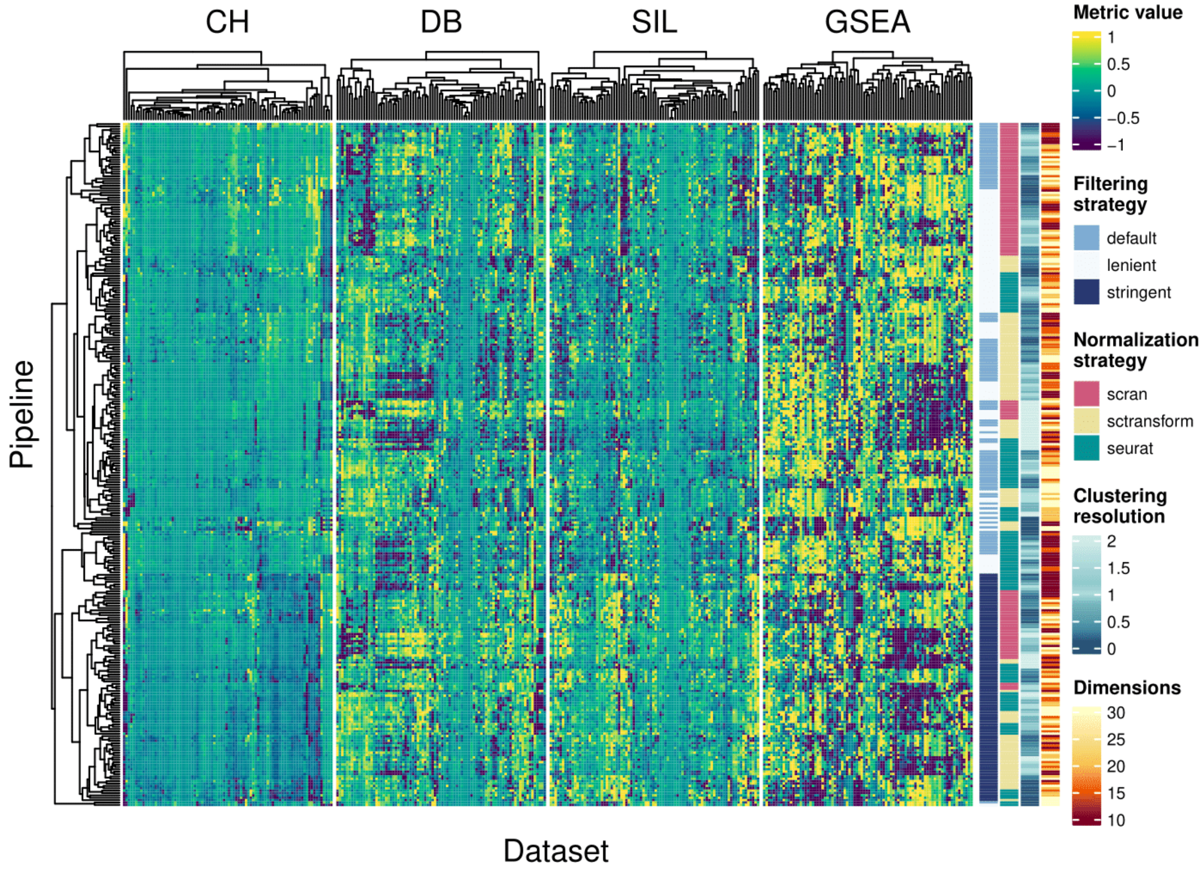


**Supplementary Figure 2** Performance of each pipeline across datasets when the effect of the number of clusters has been removed via nonparametric regression.

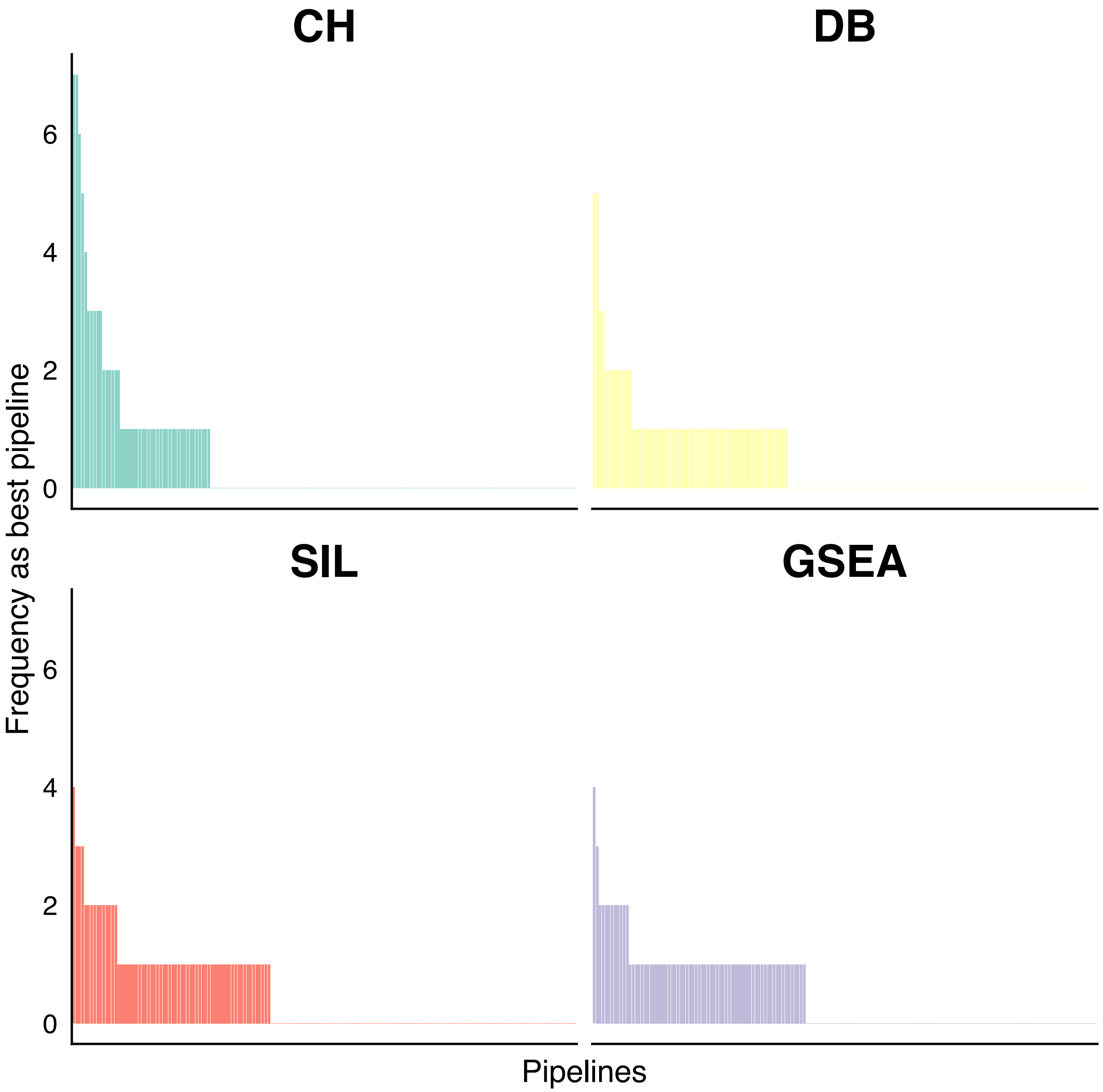


**Supplementary Figure 3** Every pipeline visualized by the number of times it had the highest metric value over all datasets. No single pipeline achieved the best performance across all datasets.

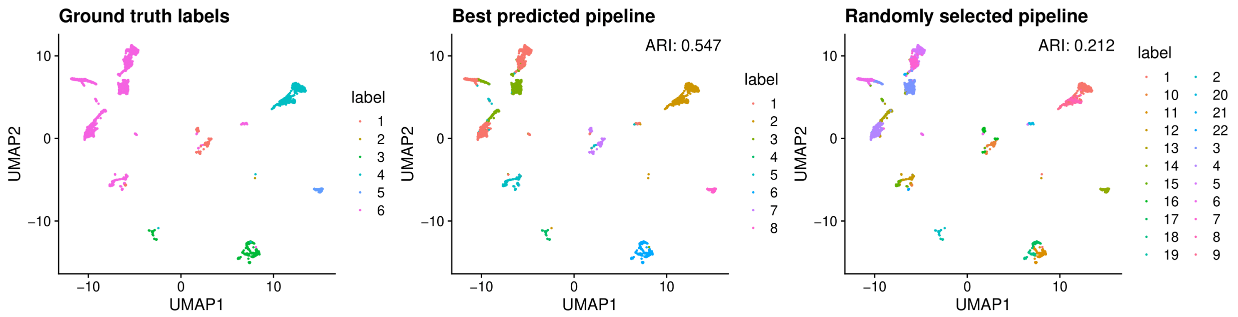


**Supplementary Figure 4** UMAP on first 50 principal components of an example dataset (EBI ID: E-MTAB-9221) with expert annotations, coloured by expert annotations (ground truth labels), labels from the best pipeline predicted by the RF with interaction terms model for SIL, and a randomly selected pipeline.
